# Vasopressin differentially modulates the excitability of rat olfactory bulb neuron subtypes

**DOI:** 10.1101/2024.06.06.597738

**Authors:** Hajime Suyama, Gaia Bianchini, Michael Lukas

## Abstract

Vasopressin (VP) is essential for social memory already at the level of the olfactory bulb (OB), and OB VP cells are activated by social interaction. However, it remains unclear how VP modulates olfactory processing to enable enhanced discrimination of very similar odors, e.g., rat body odors. So far, it has been shown that VP reduces firing rates in mitral cells (MCs) during odor presentation *in-vivo* and decreases the amplitudes of olfactory nerve-evoked excitatory postsynaptic potentials (ON-evoked EPSPs) in external tufted cells *in-vitro*. We performed whole-cell patch-clamp recordings and population Ca^2+^ imaging on acute rat OB slices. We recorded ON-evoked EPSPs as well as spontaneous inhibitory postsynaptic currents (IPSCs) from two types of projection neurons, middle tufted cells (mTCs) and MCs. VP bath-application reduced the amplitudes of ON-evoked EPSPs and the frequencies of spontaneous IPSCs in mTCs but did not change those in MCs. Therefore, we analyzed ON evoked-EPSPs in inhibitory interneurons, i.e., periglomerular cells (PGCs) and granule cells (GCs), to search for the origin of increased inhibition in mTCs. However, VP did not increase the amplitudes of evoked EPSPs in either type of interneurons. We next performed two-photon population Ca^2+^ imaging in the glomerular layer and the superficial GC layer of responses to stronger ON stimulation than during patch-clamp experiments that should evoke action potentials in the measured cells. We observed that VP application increased ON-evoked Ca^2+^ influx in juxtaglomerular cell and GC somata and decreased it in the intraglomerular neuropil. Thus, our findings indicate inhibition by VP on projection neurons via strong ON input-mediated inhibitory interneuron activity.

## Introduction

Various mammalian species rely on olfaction to identify environmental stimuli, such as food, predators, or individual conspecifics. Thus, rodents sniff conspecifics upon initiation of social behaviors and are thereby able to examine their characteristics, e.g., sex or familiarity. For example, male mice can discriminate urine from males and females even when they are not able to establish physical contact with them (Pankevich et al., 2004). Another example is social memory, also known as social discrimination (Engelmann et al., 2011), which is based on recognition of individual conspecifics encountered previously. This ability can be quantified experimentally as rats investigate an unknown stimulus rat longer than another, whom they recently had interacted with. Social memory is suggested to be highly dependent on olfaction since the olfactory bulb (OB) is essential for social discrimination (Dantzer et al., 1990).

The central actions of the neuropeptide vasopressin (VP) include the modulation of social behavior. Moreover, VP is an important enhancer of social memory (Dantzer et al., 1988). More specifically, local OB injection of a VP receptor antagonist blocks the ability to form memories of conspecifics, whereas additional local application of VP prolongs the memory for conspecifics (Dluzen et al., 1998, Tobin et al., 2010).

The OB is the very first brain region for processing and filtering olfactory signals in mammals. Neural microcircuits in the OB are known to integrate and modify signals from the olfactory epithelium before transmitting those to the olfactory cortex and other higher brain areas, which then trigger behavioral responses. Approximately 80 % of all neurons in the OB are inhibitory interneurons (Shepherd et al., 2004), such as periglomerular cells (PGCs) in the glomerular layer (GL) and granule cells (GC) in the GC layer (GCL) (Nagayama et al., 2014). Interneurons form synaptic connections onto other interneurons or projection neurons such as middle tufted cells (mTCs) and mitral cells (MCs) to inhibit them. The major portion of inhibition in the GL is suggested to function as gain control of incoming olfactory signals (e.g., Linster and Hasselmo, 1997, Cleland and Sethupathy, 2006, Cleland et al., 2007), whereas GCs organize spike timing and synchronization of projection neurons (e.g., McTavish et al., 2012, Fukunaga et al., 2014, Osinski and Kay, 2016, Egger and Kuner, 2021). Although most bulbar neurons express the classical neurotransmitters glutamate or GABA, it is known that various other substances that are released from either bulbar neurons or centrifugal projections affect neural communication in the OB as well, including neuromodulators, e.g., dopamine or acetylcholine, and neuromodulatory neuropeptides, like VP or cholecystokinin (Nagayama et al., 2014, Imamura et al., 2020, Brunert and Rothermel, 2021, Suyama et al., 2022). A source of endogenous VP that acts on social discrimination in the OB is an innate population of VP expressing cells (VPCs) that were characterized as a subpopulation of superficial tufted cells (Tobin et al., 2010, Lukas et al., 2019). As mentioned above, VP signaling at the level of the OB is essential for social memory of conspecifics (Tobin et al., 2010).

Several of our previous findings support the importance of VP neuromodulation in the OB during social memory establishment and thereby indicate intrabulbar VP release from VPCs. Thus, bulbar VPCs react with increased numbers of activated cells, i.e., positive for phosphorylated extracellular signal-regulated kinase, following social interaction *in-vivo* and with action potential firing following olfactory nerve stimulation during acetylcholine application in *in-vitro* OB slice experiments (Suyama et al., 2021). However, it is still not clear how VP modulates olfactory processing on the cellular level to enable enhanced discrimination of very similar odor mixtures, like conspecific body odors (Singer et al., 1997).

Tobin et al. (2010) showed that spontaneous firing rates and firing rates after odor stimulation in MCs decrease upon VP administration *in-vivo* and we showed that VP bath application decreases the amplitudes of electrical olfactory nerve (ON) stimulation-evoked excitatory postsynaptic potentials (EPSPs) in eTCs *in-vitro* (Lukas et al., 2019). These initial findings suggest that VP has inhibitory effects on excitatory neurons in the OB. Since the prevalent OB VP receptors are Gq-coupled excitatory V1a receptors, it is unlikely that VP acts directly on excitatory neurons. Therefore, the origin of VP inhibitory effects is not determined yet. However, we have suggestions from the morphology of VPCs. We previously showed the neurite reconstruction of VPCs including apical dendritic tufts in the GL and axons in the GCL (Lukas et al., 2019). VP-neurophysin was found in somata, dendrites, and axons, which indicates that VP could be released from dendrites, e.g., apical dendritic tufts, and axons. Biocytin-DAB reconstruction revealed that aside from apical dendritic tufts in the glomeruli, VPCs innervate densely in the GL and EPL and in the superficial GCL they have either numerous short but localized branches (type 1) or long-range projection along the internal plexiform layer (type 2) (Lukas et al., 2019). The observation leads to the hypothesis that VP is released in the GL and the superficial GCL binds to cells located there. In line with the hypothesis, strong signals of V1a-receptor staining were observed in the GL (Ostrowski et al., 1994) and in the superficial part of the GCL (Tobin et al., 2010). Moreover, discriminability of similar odors including social discrimination is regulated by bulbar interneurons, which in turn modulate excitatory/projection neurons (Abraham et al., 2010, Oettl et al., 2016). Thus, we hypothesized that an excitatory action of VP on inhibitory neurons in the GL or the GCL enhances inhibition of excitatory neurons during social interactions.

As a first step to examine how VP might enhance the inhibition of excitatory neurons during odor processing, we investigated VP effects on the responses of different cell types, including both, excitatory and inhibitory neurons, to ON stimulation in acute OB slices, which mimics sensory activation. Therefore, we recorded ON-evoked EPSPs and spontaneous inhibitory postsynaptic currents (IPSCs) in mTCs and MCs that project to the cortices, and ON-evoked EPSPs in PGCs and GCs. Furthermore, we performed Ca^2+^ population imaging of ON-evoked responses in the GL and the superficial GCL using two-photon microscopy.

## Materials and Methods

### Animals

All experiments were conducted according to guidelines for the care and use of laboratory animals of the local government of Oberpfalz and Unterfranken. Wistar rats of either sex were purchased from Charles River Laboratories (Sulzfeld, Germany) or bred onsite in the animal facilities at University of Regensburg. The light in the rooms was set to an automatic 12 h-cycle (lights on 07:00-19:00)

### Slice preparation

11-18 day-old juvenile rats of either sex were used for *in-vitro* electrophysiology and Ca^2+^ imaging experiments. The rats were deeply anesthetized with isoflurane and quickly decapitated. Horizontal slices (300 µm) of the OB were cut in ice-cold, carbogenized ACSF (artificial cerebrospinal fluid; in mM: 125 NaCl, 26 NaHCO3, 1.25 NaH2PO4, 20 glucose, 2.5 KCl, 1 MgCl, and 2 CaCl2) using a vibratome (VT 1200, LEICA, Wetzlar, Germany) and afterwards incubated in ACSF at 36 °C for 45 min. Until experiments, the slices were kept at room temperature (∼21 °C) in ACSF.

### Electrophysiology

Brain slices were placed in a recording chamber on the microscope’s stage perfused with carbogenized ACSF circulated by a perfusion pump (ISM 850, Cole-Parmer, Wertheim, Germany). To perform whole-cell patch-clamp recordings, cells were visualized by infrared gradient-contrast illumination via an IR filter (Hoya, Tokyo, Japan). Glass pipettes for recordings were pulled by a pipette puller (Narishige, Tokyo, Japan) sized 4-6 MΩ and filled with intracellular solution. The intracellular solution for current-clamp recordings contained 130 K-methylsulfate, 10 HEPES, 4 MgCl2, 4 Na2ATP, 0.4 NaGTP, 10 Na Phosphocreatine, and 2 ascorbate (in mM) at pH 7.2, and the intracellular solution for voltage-clamp recordings contained 110 CsCl, 10 HEPES, 4 MgCl2, 10 TEA, 10 QX-314, 2.5 Na2ATP, 0.4 NaGTP, 10 Na Phosphocreatine, and 2 ascorbate (in mM) at pH 7.2. Recordings were performed with an EPC-10 (HEKA, Lambrecht, Germany) digital oscilloscope. Series resistance was ranging between 10 to 30 MΩ. The average resting membrane potentials were -60 to -70 mV in MCs and mTCs, -50 to -60 mV in PGCs, and -70 to -80 mV in GCs. Experiments were only started in case the patched cells had a holding current below approximately -50 pA and a stable resting membrane potential. Experiments were performed at room temperature (∼21 °C). ON stimulation was performed with a glass pipette stimulation electrode sized around 2 MΩ. Glass pipettes were filled with ACSF. The unipolar electrode was connected to an external stimulator (STG 1004, Multi-Channel Systems, Reutlingen, Germany). The stimulation strength was adjusted via the stimulator’s software (MC_Stimulus, v 2.1.5) and stimulation was triggered by the amplifier software (Patchmaster, v2×73.5, HEKA). Stimulation pipettes were gently placed in the ON layer anterior to the area selected for patching using a manual manipulator (LBM-7, Scientifica, East Sussex, UK) under optical control with the microscope. The stimulation lasted for 100 µs, with a current of 20-200 µA for mTCs, 25-350 µA for MCs, 9-100 µA for PGCs, or 5-150 µA for GCs. We confirmed the identity of PGCs with post-hoc morphological examination. We added biocytin (5 mg/mL, Sigma-Aldrich, Darmstadt, Germany) in the intracellular solution to fill cells during recording and subsequently visualized apical dendrite arbors using enzymatic 3,3’-Diaminobenzidine based staining (Vector Laboratories, CA, US) (Lukas et al., 2019). All patched putative PGCs had a soma sized <10 µm and no long-range laterally projecting neurite, confirming their identity as PGCs (Nagayama et al., 2014).

#### Experimental design

In current-clamp experiments recording ON-evoked EPSPs, ON stimulation was triggered only every 30 s to prevent run-down (Lukas et al., 2019). VP was diluted in ACSF ([Arg^8^]-vasopressin acetate salt, Sigma-Aldrich, Darmstadt, Germany, 1 µM) and bath-applied via the perfusing system after a baseline recording of 5 min. Traces in the VP condition were recorded no earlier than 5 min after the onset of administration. Traces were averaged over 5 stimulations and two such averaged traces each in the ACSF condition and in the VP condition were analyzed. Averaged amplitudes within conditions were normalized to the ACSF condition (100%). Data was analyzed with Origin 2020 (Origin Lab Corporation, Northampton, MA, US).

In voltage-clamp experiments spontaneous IPSCs were recorded at 0 mV for 10 min during each condition. VP (1 µM) was bath-applied via the perfusion system and the VP condition was recorded 5 min after the onset of administration for 5 min. Frequencies and amplitudes of IPSCs were normalized to the ACSF condition (100%). Data was analyzed with the Peak Analyzer in Origin 2020.

### Population Ca^2+^ imaging

For population Ca^2+^ imaging, the AM-dye Cal 520 (1 µM, AAT Bioquest, CA, USA) and Alexa 594 (50 µM, Invitrogen) were loaded into the superficial GCL or the GL via a glass pipette sized around 2 MΩ. Loading pipettes were guided by light microscopy and by the Alexa 594 fluorescence. The Ca^2+^ dye was loaded for 15 s using the picospritzer III device (Parker Hannifin, NH, USA) followed by 20-min incubation to let the Ca^2+^ dye be taken up by cells. The fluorescence was imaged at a wavelength of 850 nm in raster-scan mode using a two-photon resonant scanner (frame rate of 31.5 Hz). Femto-2D microscope (Femtonics) was equipped with a Mai Tai wideband, mode-locked Ti:Sapphire laser (Spectra-Physics, CA, USA) and a 20x Zeiss water-immersion objective (Carl Zeiss, Oberkochen, Germany). The microscope was controlled by MESc v3.3.4290 software (Femtonics). ON stimulation (400 µA, 100 µs) was applied three times for each condition, control (ACSF) and VP (1 µM). VP was bath-applied via the perfusion system and the VP condition was recorded 10 min after the onset of administration.

The raw data of the experiments was imported to Fiji (ImageJ, downloaded from https://imagej.net/Fiji/Downloads) and ΔF/F in the somata of GCs, juxtaglomerular cells (JGCs, decided by the small cell bodies), and the glomeruli was extracted using the ROI selection tool. The resulting traces from the 3 stimulations per condition were averaged. ΔF/F amplitudes were analyzed with Origin 2020. Averaged amplitudes within conditions were normalized to the ACSF condition (100%).

Ca^2+^ imaging experiments were performed at room temperature (∼21 °C).

### Statistics

Statistics were performed with SPSS (ver. 26, IBM, Armonk, NY, USA). All statistical analysis performed was two-sided and significance was accepted at p<0.05. All data in the text are shown with average ± standard deviation.

## Results

### VP reduced ON-mediated excitation and increased spontaneous inhibition in mTCs but not in MCs

We performed patch-clamp recordings in either current-clamp or voltage-clamp configuration in mTCs and MCs in acute OB slices (Figure 1A). Electrical ON stimulation reliably evoked EPSPs in mTCs and MCs. We then compared EPSP amplitudes in the presence of VP (1 µM in ACSF) to the control condition (ACSF). In mTCs VP reduced the amplitudes of ON-evoked EPSPs to 60.4 ± 20.5 % of control (p=0.012, z=-2.521, related samples Wilcoxon test, n=8 from 6 rats. Figure 1B,C). The amplitudes of ON-evoked EPSPs without VP application were stable over time compared to the VP condition (95.0 ± 4.6 % of control, 10 min after the start of the measurement. n=5 from 4 rats. p=0.004, t(11)=3.657, t-Test vs. VP). Therefore, we concluded that reduction is due to VP application but not desensitization of bulbar circuits to ON stimulation. In another set of experiments using voltage-clamp recordings, VP increased the frequencies of spontaneous IPSCs to 123.3 ± 22.1 % of control (p=0.012, z=2.521, related samples Wilcoxon test, n=8 from 4 rats. Figure 1D,E). However, the amplitudes of spontaneous IPSCs were not changed (106.9 ± 22.7 %. p=0.263, z=1.120, related samples Wilcoxon test, n=8 from 4 rats. Figure 1F), indicating a predominantly presynaptic effect. Furthermore, in all experiments (n=8 from 4 rats) spontaneous IPSCs were abolished following bath application of bicuculline (a GABA receptor antagonist, 50 µM) confirming the GABAergic origin of these signals (Data not shown). Therefore, the data implies that VP enhances both ON-evoked and tonic inhibitory modulation of mTCs. Surprisingly, we did not observe any of those VP inhibitory effects in MCs, even though broad distributions in evoked EPSP amplitudes and IPSC frequencies in the VP condition were observed (ON-evoked EPSPs: 99.4 ± 20.2 % of control. p=0.929, z=-0.089, related samples Wilcoxon test, n=11 from 9 rats. Figure 1G, H; Frequencies of spontaneous IPSCs: 105.3 ± 38.2 % of control. p=0.889, z=0.140, related samples Wilcoxon test. n=8 from 6 rats. Figure 1I, J; Amplitudes of spontaneous IPSCs: 97.2 ± 10.6 % of control. p=0.779, z=-0.280, related samples Wilcoxon test, n=8 from 6 rats. Figure 1I, K). This lack of consistent inhibitory effects in MCs was somewhat unexpected, as Tobin et al. (2010) showed *in-vivo* that VP reduces spontaneous and odor-evoked spiking rates in MCs.

**Figure 1.**
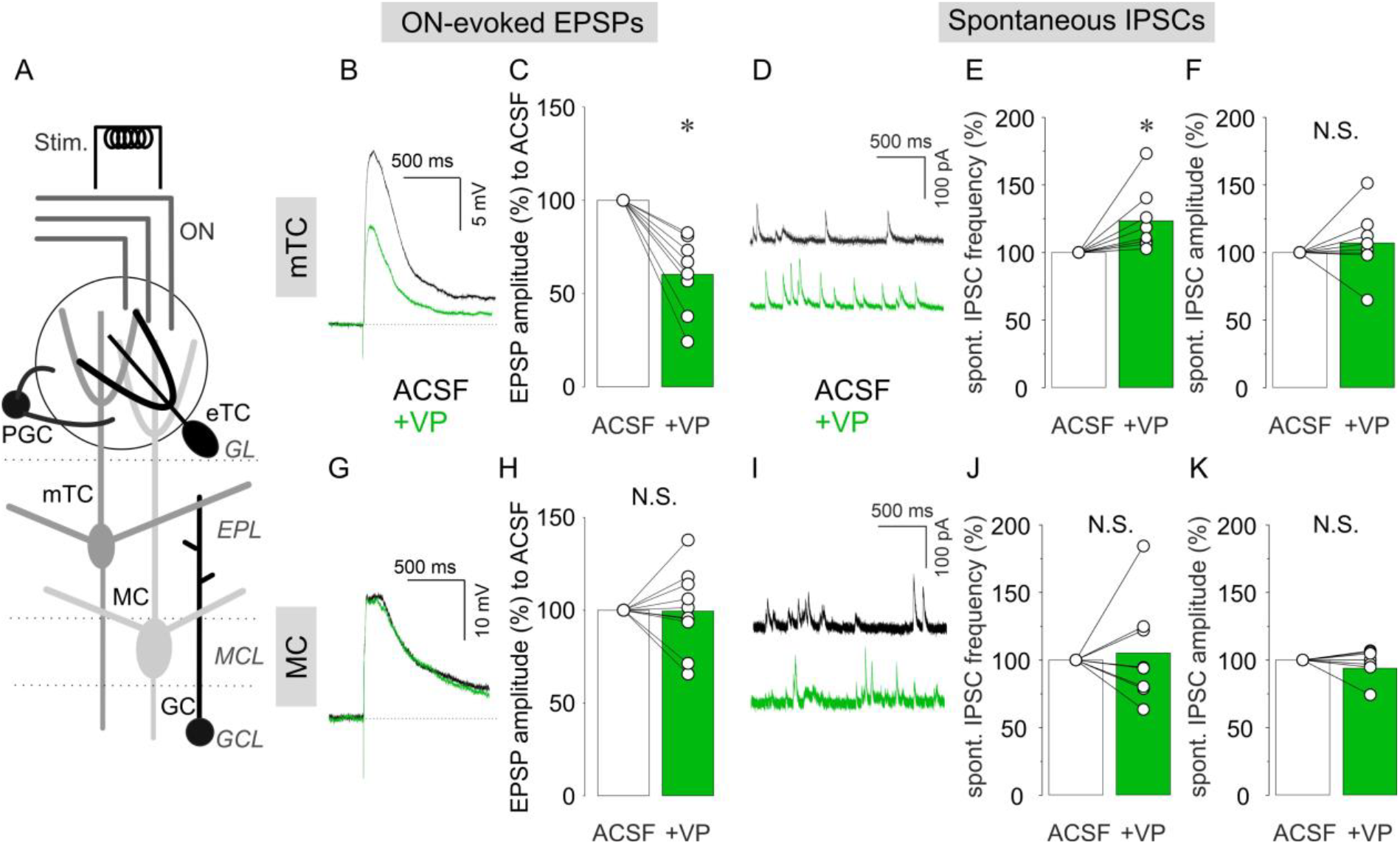
Patch-clamp recordings from mTCs and MCs. (A) the schematic image of the OB slice. ON, olfactory nerve: GL, glomerular layer: EPL, external plexiform layer: MCL, mitral cell layer: GCL, granule cell layer: Stim., electrical ON stimulation: PGC, periglomerular cell: eTC. external tufted cell: mTC, middle tufted cell: MC, mitral cell: GC, granule cell. Representative current-clamp traces of ON-evoked EPSPs and cumulative data of ON-evoked EPSP amplitudes in % to control in mTCs (B, C) and MCs (G, H). Representative voltage-clamp traces of spontaneous IPSCs and cumulative data of spontaneous IPSC frequencies and amplitudes in % to control in mTCs (D, E, F) and MCs (I, J, K). Bar graphs show average values. Individual data points are shown as open circles and points from the same cell are connected by a line. *, p<0.05: N.S., not significant. Black traces, ACSF (control): Green traces, VP.

### VP did not increase evoked EPSPs in inhibitory neurons

Where does the inhibition onto excitatory projection neurons, e.g., eTCs or mTCs, originate from? There are two main populations of inhibitory interneurons in the OB. One is in the glomerular layer (GL), where synapses between the ON and bulbar neurons reside within the glomerular neuropil. Thus, a first modulation of olfactory inputs takes place in this layer. There are many cell types of inhibitory neurons in the GL and the most numerous are PGCs (Nagayama et al., 2014). We performed patch-clamp recordings in PGCs regardless of subtypes, as we were not able to differentiate them in our experimental setup. ON stimulation evoked EPSPs in PGCs. We observed mixed effects of VP (1 µM) on the amplitudes of evoked EPSPs including increase, decrease, or no changes (Figure 2A). Consequently, there was no overall significant difference between VP application compared to the control condition (85.4 ± 26.1 % of control. p=0.131, z=-1.511, paired Wilcoxon test, n=11 from 9 rats. Figure 2B). We further categorized PGCs into either Type A or Type C according to their firing patterns as described by Tavakoli et al. (2018). In addition, we examined if hyperpolarizing currents evoked sags due to hyperpolarization-induced depolarization. Also, we visualized patched putative PGCs by filling cells with biocytin (see methods) to investigate their morphology. However, we could not find any correlation between these electrophysiological or morphological characteristics and the different directions of VP effects (data not shown).

**Figure 2.**
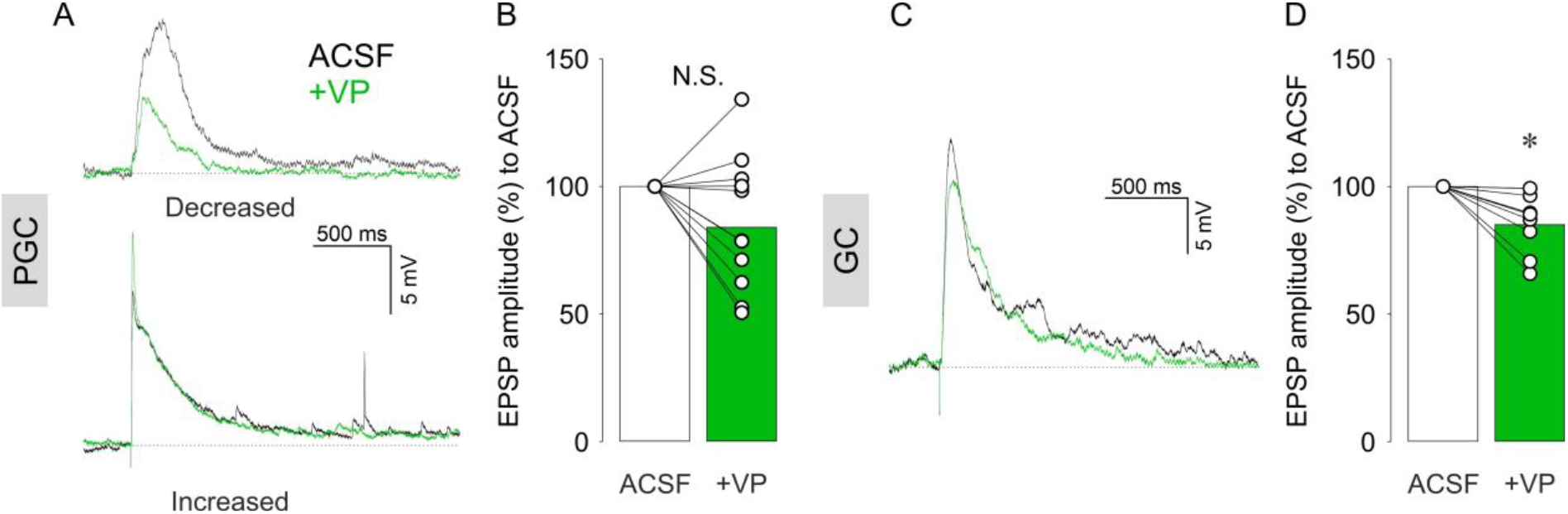
Representative current-clamp traces of ON-evoked EPSPs and cumulative data of ON-evoked EPSP amplitudes in % to control in PGCs (A,B) and GCs (C,D). Bar graphs show average values. Individual data points are shown as open circles and points from the same cell are connected by a line. *, p<0.05: N.S., not significant. Black traces, ACSF(control): Green traces, VP.

We, next, examined the second main population of inhibitory interneurons in the OB, the GCs (Nagayama et al., 2014). Unlike PGCs, VP consistently decreased the amplitudes of ON-evoked EPSPs in GCs to 85.0 ± 11.7 % of control (p=0.012, z=-2.521, related samples Wilcoxon test, n=8 from 8 rats. Figure 2C, D). Thus, ON-evoked EPSPs were not increased upon VP administration in both inhibitory interneuron populations, arguing against an increased excitation of interneurons via the sensory afferents as mechanism for the inhibitory action of VP on eTCs and mTCs.

### VP increased evoked Ca^2+^ influx in inhibitory neurons

While VP did not increase the subthreshold excitability of inhibitory neurons, we next wondered whether suprathreshold activation might be enhanced. Therefore, we performed two-photon population Ca^2+^ imaging to examine VP effects on Ca^2+^ influx in inhibitory interneurons with strong ON stimulation which is likely to evoke action potentials in stimulated cells from our experience (400 µA: up to 100 µA or 150 µA for EPSP experiments in PGCs or GCs, respectively). We injected the AM-dye Cal-520 into the GL followed by two-photon imaging with a resonant scanner. After JGCs took up the dye into their somata, we stimulated the ON and measured ΔF/F in JGC somata as well as in the neuropil in the glomeruli (Figure 3A, B, Egger et al., 2003). While there were mixed effects, on average VP significantly increased the amplitudes but not the integral of ON-evoked ΔF/F changes to 117 ± 52.5 % and 109 ± 69.3 % of control, respectively (Amplitudes, p=0.001, z=3.193, related samples Wilcoxon test; Integral, p=0.619, z=-0.497, related samples Wilcoxon test. n=166 from 5 rats. Figure 3B, C). In contrast, the amplitudes and the integral of ON-evoked ΔF/F in the glomeruli were significantly decreased to 91.9 ± 9.1 % and 84.0 ± 19.9 % of control, respectively (Amplitudes, p=0.015, z=-2.432, related samples Wilcoxon test; Integral, p=0.019, z=-2.353, related samples Wilcoxon test. n=12 from 5 rats. Figure 3B, D). The neuropil in the glomeruli consists of ON axons, apical dendritic tufts of OB projection neurons and neurites of JGCs. Therefore, the reduction of ON-evoked Ca^2+^ influx in the glomeruli by VP might reflect the increasing effects of VP on JGCs that are predominantly inhibitory neurons and thus can be expected to inhibit the intraglomerular neuropil more strongly. We also performed dye injections in the superficial GCL and measured changes in intracellular Ca^2+^ levels in GC somata (Figure 4E, F). In these experiments, again there were mixed effects but on average VP significantly increased the amplitudes and the integral of ON-evoked ΔF/F changes to 128 ± 56.6 % and 176 ± 153 % of control, respectively (Amplitudes: p<0.001, z=5.912, related samples Wilcoxon test; Integral: p<0.001, z=7.243, related samples Wilcoxon test, n=165 from 6 rats. Figure 3G, H).

**Figure 3.**
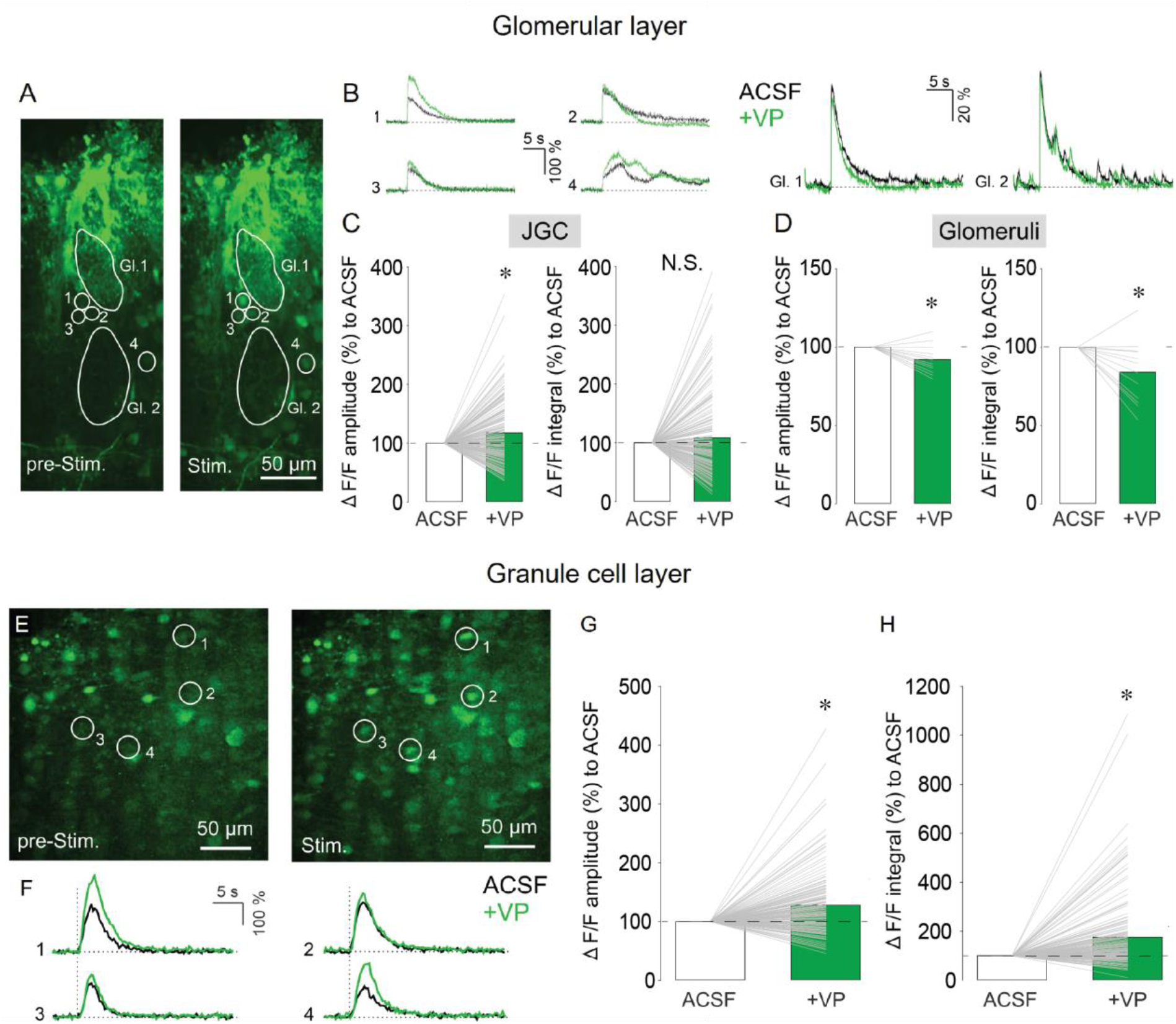
Representative images of Ca2+ imaging before ON stimulation (left) and after ON stimulation (right) in the GL (A). Gl., glomerulus. Representative traces of Ca2+ imaging from corresponding JGCs (left) and from corresponding glomeruli (right) in A. Cumulative data of amplitudes (left) and integral (right) of ΔF/F in JGCs (C) and glomeruli (D). Bar graphs show average values. Individual data points from the same cell are connected by a line. *, p<0.05. Black traces, ACSF (control): Green traces, VP. Representative images of Ca2+ imaging before ON stimulation (left) and after ON stimulation (right) in the GCL (E). Representative traces of Ca2+ imaging from corresponding GCs in E (F). Cumulative data of amplitudes (G) and integral (H) of ΔF/F in GCs. Bar graphs show average values. Individual data points from the same cell are connected by a line. *, p<0.05. Black traces, ACSF (control): Green traces, VP.

**Figure 4.**
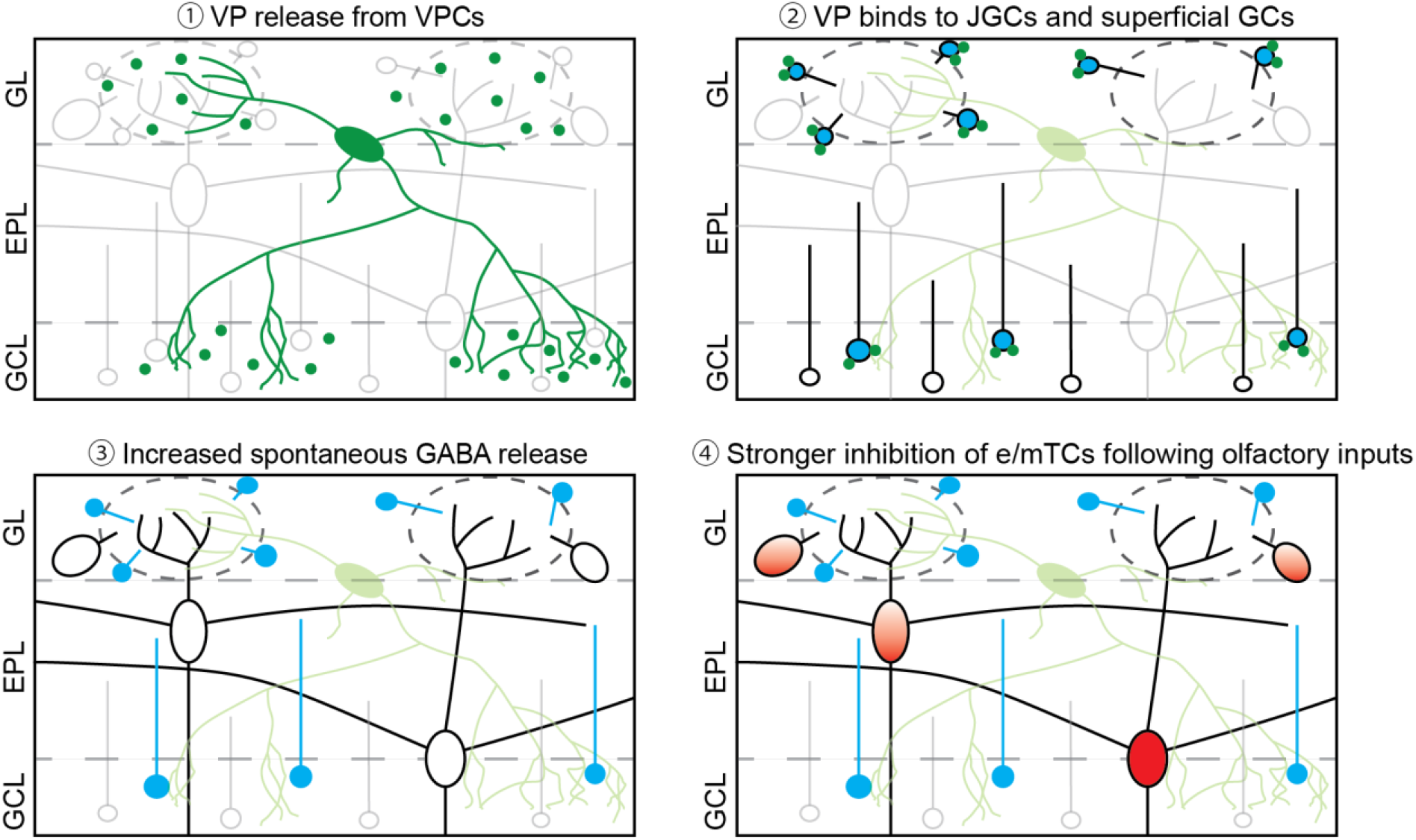
Scheme of the overview 1, Once VPCs are activated, VP is released from their dendrites and axons in both the GL and the superficial GCL. 2, Released VP binds to VP receptors, which are expressed in the GL and the superficial GCL, likely on JGCs and GCs. 3, By VP receptor activation, spontaneous GABA release occurs more frequently leading to more spontaneous inhibitory inputs onto mTCs. 4, Following olfactory inputs, VP-bound interneurons are activated stronger and then inhibit e/mTCs.

Therefore, we suggest that VP enhances evoked Ca^2+^ influx in those inhibitory interneurons, which in turn might increase both the spontaneous and evoked GABA release probability onto mTCs.

## Discussion

### Origin of VP inhibitory effects in the OB

V1a receptors that are predominantly expressed in the olfactory bulb are Gαq/11-coupled receptors and thus act in an excitatory manner (Birnbaumer, 2002). As mentioned in the introduction, V1a receptors are expressed in the GL and the superficial part of the GCL (Ostrowski et al., 1994, Tobin et al., 2010). Also, VPCs innervate densely those two layers in the OB (Figure 4)(Figure 4, Lukas et al., 2019). This distribution of V1a receptors and VPC’s innervation fit our data showing that VP increased the ON-evoked Ca^2+^ signal in JGCs and GCs in those layers and decreased the evoked Ca^2+^ signal in the glomeruli. Moreover, VP increases inhibition of eTCs (Lukas et al., 2019) and mTCs (Figure 4). Similar VP-mediated increase of inhibition has been shown in other brain regions. For example, VP increases the frequency of spontaneous IPSP/Cs in magnocellular paraventricular nucleus neurons (Hermes et al., 2000), as well as spontaneous spikes in hippocampal GABAergic neurons which results in an increase in the amount of spontaneous IPSCs in pyramidal neurons (Ramanathan et al., 2012). Ramanathan et al. (2012) suggested that increased hippocampal GABAergic inhibition onto pyramidal cells may result in organized inhibition to establish fine tuning of excitation and rhythmic synchronized activity of pyramidal neurons that are important for memory consolidation. Since the function of bulbar inhibitory networks is suggested to tune excitation and modulate oscillatory activity as well, VP effects on bulbar inhibitory neurons may help organization of excitatory neuron activity.

Functional differences between PGCs (inhibition in the GL) and GCs have been discussed extensively elsewhere (see review, Devore and Linster, 2012, D’Souza and Vijayaraghavan, 2014). Briefly, inhibition in the GL tunes activation patterns of projection neurons, such as contrast enhancement or concentration invariance (e.g., Linster and Hasselmo, 1997, Cleland and Sethupathy, 2006, Cleland et al., 2007). For instance, acetylcholine (ACh) which is known to activate GL interneurons (Devore and Linster, 2012), enhances contrast of projection neural representation responding to very similar odors. Thus, neostigmine (an acetylcholinesterase inhibitor) administration in the OB increases differences in number of MC spikes between responses to ethers differing by single carbon chain in *in-vivo* electrophysiology. For example, a MC responds strongest to E2 (ethyl acetate) and second strongest to E3 (ethyl propionate), differing by only one carbon chain, but the responses are not significantly different each other. At the presence of neostigmine, the MC responds less strongly to, or gets more inhibited by E3 than control. Therefore, responses of the MC to E2 and E3 are discriminable (Chaudhury et al., 2009). Moreover, electrical stimulation of the horizontal limb of the diagonal band of Broca (HDB), the center of cholinergic top-down projections, decreases glomerular M/TC-tuft activity following odor presentation with high concentration whereas increases glomerular activity following low concentration odor presentation in *in-vivo* Ca^2+^ imaging, indicating responses are less varied to different concentrations of the same odor (Bendahmane et al., 2016). These authors suggested that the less variation of responses to different odor concentrations makes neural representation reflect more purely the identity of odors.

GCs are responsible for organization of spike timing and synchronization of projection neurons that further refines contrast and representations of odors (e.g. McTavish et al., 2012, Osinski and Kay, 2016). Computational analysis showed that the level of GC excitability might tune oscillatory frequency in MCs, thus with low excitability of GCs, MCs fire in the gamma range, however high excitability allows MCs to fire in beta oscillation (Osinski and Kay, 2016). Oettl et al. (2016) showed the modulation of GCs via the oxytocin system in the anterior olfactory nucleus (AON). An oxytocin receptor agonist increases the frequency of spontaneous EPSCs in GCs from AON excitatory neurons resulting in increased spontaneous IPSCs in MCs. In *in-vivo* electrophysiological recordings in M/TCs, an oxytocin receptor agonist applied in the AON lowers basal spiking rates and increases odor-evoked spiking rates, thus improve the signal-to-noise ratio. Conditional oxytocin receptor-knockout in the AON impairs social memory showing that enhanced signal-to-noise in M/TCs by activation of GCs is important for discrimination of conspecific body odors (Oettl et al., 2016). Moreover, we observed that VP application increased the amplitudes of ON-evoked Ca^2+^ influx into GCs in population imaging, which is an opposite effect to VP action on ON-evoked EPSPs. The possible explanation would be the different ON stimulation intensities, causing subthreshold versus suprathreshold activation of GCs. Apparently, VP can enhance Ca^2+^ entry selectively for suprathreshold activation, which is probably not related to the initial depolarization but to later phases of the signal which involve contributions from NMDARs, both of which are present at GC somata (Personal communication with M Sassoe-Pognetto,Stroh et al., 2012). In rat ventral hippocampus, VP enhances glutamate-evoked spiking, and the VP effects are blocked by both V1a receptor antagonist and NMDAR antagonist, indicating VP effects via modulation of NMDARs (Urban and Killian, 1990). We have not performed experiments on interneurons in the EPL (Nagayama et al., 2014). Since VPC’s neurites are found through the EPL, it would be informative to investigate VP effects on them as well.

Our data showed that VP reduced the amplitudes of ON-evoked EPSPs in eTCs (Lukas et al., 2019) and mTCs. These data indicate that eTCs and mTCs need stronger ON inputs to fire under the VP condition. The suppression of those neurons could result in contrast enhancement of the neural representation in the OB (Chaudhury et al., 2009). The glomerular GABAergic inhibition includes GABAA receptor- and GABAB receptor-mediated pathways. For instance, cholecystokinin which is also expressed in a subpopulation of superficial tufted cells, acts on SACs to inhibit presynaptically the ON via GABAB receptors resulting in smaller ON-evoked EPSCs in eTCs (Liu and Liu, 2018). Therefore, a similar mechanism is conceivable for VP modulation and further examination of the involvement of SACs or GABAB receptors would give us more insights in the VP-mediated inhibition of eTCs and mTCs. Interestingly, unlike for mTCs or eTCs, VP did not reduce the amplitudes of evoked EPSPs or frequencies of spontaneous IPSCs in MCs. This variance in our data between mTCs and MCs could be due to the difference on input sources of the two cell types. MCs receive indirect excitatory inputs from eTCs (De Saint Jan et al., 2009, Najac et al., 2011), and MCs are less sensitive to odor inputs than e/mTCs (Igarashi et al., 2012). If the ON-evoked EPSPs in MCs are consequences of firing of eTCs, the ON stimulation intensity in MC experiments may be strong for eTCs, so that also during VP application still APs are evoked in eTCs. Thus, during VP application the same net amount of excitation is transmitted to MCs as without VP. In addition, since we did not perform Ca^2+^ imaging in MCs with evoked APs, we cannot exclude that VP inhibitory effects in MCs may be observed when they fire. Therefore, Ca^2+^ imaging might show similar inhibitory effects in firing MCs and would then be in line with Tobin et al. (2010) that showed a decrease in spiking rates *in-vivo* following topical VP application. Since Tobin et al. (2010) reported that V1a receptors are expressed in MCs as well, we cannot exclude that VP directly excites MCs, even though we could not see excitatory effects in our experiments. In CA1, VP increases not only the number of spontaneous IPSCs but also the number of spontaneous spikes under the condition with glutamatergic and GABAergic receptor antagonists in pyramidal neurons (Ramanathan et al., 2012). Therefore, the non-synaptic excitability of projection neurons would be intriguing to examine.

### Possible consequences of VP-mediated OB modulation

We previously suggested that social discrimination is a variation of perceptual learning because of the association with ACh and close similarity of stimuli, i.e., conspecific body odors, that rats discriminate (Suyama et al., 2021). Perceptual learning is possible, because a subject pays attention to a stimulus during exposures then sensory acuity against the stimulus is enhanced due to a finer neural representation. As mentioned above, ACh, an important substance for perceptual learning, increases differences in numbers of MC spikes reacted to odors differing by a single carbon (Chaudhury et al., 2009). This change in neural representation results in behavioral outputs, like habituation. Rats lose their motivation to investigate odor if rats perceive it as the same one as they previously investigated, i.e., habituation. In controls, rats show habituation to an odor differing by a single carbon from a previously exposed odor suggesting that rats cannot distinguish odors differing by one carbon. However, injection of neostigmine into the OB enables rats to discriminate those odors (Chaudhury et al., 2009). Therefore, neuromodulation of projection neuron activity seems to be important for sensory perception, hence discrimination as behavioral output of this enhanced sensory perception. We demonstrated that VP modulation may result in less mTCs firing via GL contrast enhancement. Further, VP inhibits firing rates of MCs (Tobin et al., 2010) which can result in an improved neural representation (Linster and Hasselmo, 1997). Although Tobin et al. (Tobin et al., 2010) did not record the firing rates of mTCs, it is plausible that this is the same for all projection neurons. Thus, in the VP condition M/TCs may transmit more precise information to higher brain regions, like during the modulatory action of ACh (Chaudhury et al., 2009). Taken together, we hypothesize that VP inhibits projection neurons differently to reduce sensitivity and improve representation in the olfactory cortex.

## Ethics statement

All experiments were conducted according to national and institutional guidelines for the care and use of laboratory animals, the rules laid down by the EC Council Directive (86/89/ECC) and German animal welfare.

## Author contributions

H.S. and M.L. designed the research and performed patch-clamp experiments and analysis. G.B. and M.L. performed Ca^2+^ population imaging and analysis. H.S., and M.L. wrote and revised the manuscript.

## Acknowledgements

We would like to thank Anne Pietryga-Krieger for experimental support, Christoph Schmid and Atefeh Akbari for help with experimentation, and Veronica Egger for experimental equipment such as electrophysiology rigs and Ca^2+^ imaging setups and for advice.

